# Structural insights into the force-transducing mechanism of a motor-stator complex important for bacterial outer membrane lipid homeostasis

**DOI:** 10.1101/2024.11.27.625625

**Authors:** Jiang Yeow, Chee Geng Chia, Nadege Zi-Lin Lim, Shu-Sin Chng

**Affiliations:** Department of Chemistry, Faculty of Science, National University of Singapore, Singapore 117543; National University of Singapore Graduate School for Integrative Sciences and Engineering, Singapore 117456; Singapore Center for Environmental Life Sciences Engineering, National University of Singapore (SCELSE-NUS), Singapore 117456

**Keywords:** Cryo-EM, proton motive force, rotary motion, conformational changes, TolQRA

## Abstract

Gram-negative bacteria assemble an asymmetric outer membrane (OM) that functions as an effective barrier against antibiotics. Building a stable and functional OM requires assembly and maintenance of balanced levels of proteins, lipopolysaccharides, and phospholipids into the bilayer. In *Escherichia coli*, the trans-envelope Tol-Pal complex has recently been established to play a primary role in maintaining OM lipid homeostasis. It is believed that the motor-stator complex TolQR exploits the proton motive force in the inner membrane to induce conformational changes in the TolA effector, ultimately generating a force across the cell envelope to activate processes at the OM. Molecular details of how such force transduction occurs via the TolQRA complex is unknown. Here, we solve structures of the *E. coli* TolQRA complex using single particle cryo-EM, capturing the transmembrane (TM) regions of the purified complex in two distinct states at ∼3.6 Å and ∼4.2 Å nominal resolutions. We define how the TolA N-terminal TM helix interacts with an asymmetric TolQ_5_R_2_ sub-complex in two different positions, revealing how the two TolQRA states are related by rotation of the TolQ pentamer. By considering structural prediction of the periplasmic domains of the complex, we propose a working model for how proton passage through the complex induces rotary movement that can be coupled to TolA for force transduction across the cell envelope.

## Introduction

A defining characteristic of the Gram-negative bacterial cell envelope is the presence of the outer membrane (OM), an asymmetric lipid bilayer that is crucial for growth and survival (Nikaido, 2003). The OM is assembled ∼210±27 Å from the cytoplasmic or inner membrane (IM), creating an envelope structure that encompasses an aqueous periplasmic compartment also containing the bacterial cell wall (Matias et al., 2003). While this multi-layered architecture offers protection against external threats such as antibiotics, it also poses huge challenges for molecular transport, especially in the context of building and maintaining the OM (Guest & Silhavy, 2023), as well as acquiring scarce but essential nutrients for growth (Nikaido, 2003). In particular, the periplasm and OM are devoid of typical energy sources such as ATP, thereby necessitating specialized mechanisms to enable energetically demanding processes to occur within the cell envelope.

One such mechanism is the transduction of the proton motive force (*pmf*) at the IM to a physical force across the cell envelope, via the so-called motor-stator complexes. These complexes are important for powering motility (e.g. MotAB and AglRQS (Sridhar, 2020; Sun et al., 2011)), metal-siderophore uptake (e.g. ExbBD (Ratliff et al., 2022)), or OM lipid homeostasis (e.g. TolQR (Shrivastava et al., 2017; Szczepaniak et al., 2020; Tan & Chng, 2024)). Structural studies have revealed a consistent 5:2 architecture of the motor-stator complexes (MotA_5_B_2_ (Deme et al., 2020a; Santiveri et al., 2020), ExbB_5_D_2_ (Celia et al., 2019; Maki-Yonekura et al., 2018), TolQ_5_R_2_ (Karimullina et al., 2024; Williams-Jones et al., 2023)), where a pentameric motor forms a ring of transmembrane (TM) helices that encapsulates a pair of N-terminal helices of the dimeric stator. Each motor subunit comprises at least three membrane-spanning helices, while each stator contains a periplasmic domain linked to a single TM helix. Missing from the solved structures, however, the dimeric periplasmic domains of the stator are known to bind the cell wall, thereby anchoring the complexes in place (Roujeinikova, 2008; Wojdyla et al., 2015; Zinke et al., 2023). Proton translocation across these 5:2 motor-stator complexes, down its electrochemical gradient, is thought to drive rotation of the motor by changing the protonation states of conserved aspartate residues in the pair of TM helices of the stator (Blair & Berg, 1988; Braun & Herrmann, 1993; Cascales et al., 2001). In motility, such rotation in multiple MotA_5_B_2_ complexes generates the massive torque required to turn the flagellum, which propels the bacterium forward (Johnson et al., 2024; Singh et al., 2024).

How ExbBD and TolQR motor-stator complexes utilize rotary movement for force transduction is less clear (Ratliff et al., 2022; Szczepaniak et al., 2020). Interestingly, both complexes are associated with an additional ‘effector’ protein, TonB and TolA, respectively, that is believed to somehow convert motor rotation in the plane of the IM into an orthogonal force applied across the cell envelope. Both effector proteins comprise an N-terminal TM helix (domain I) that interacts with the motor-stator in the IM, and a structurally-homologous C-terminal periplasmic domain (domain III) that engages OM components for corresponding function (Levengood et al., 1991; Postle & Good, 1983; Sean Peacock et al., 2005). The key difference between TonB and TolA lies in the linker domain (domain II) – TonB-II is proline-rich and possibly less structured (Brewer et al., 1990; Domingo Köhler et al., 2010), while TolA-II contains repeats of an alanine-rich heptad sequence that may favor helical structures (Witty, 2002). These architectural differences imply distinct mechanisms of trans-envelope force transmission by the respective complexes, perhaps also consistent with their separately evolved functions. In the case of ExbBD-TonB, force transduction triggers metal-siderophore passage across TonB-dependent transporters (TBDTs) in the OM upon interaction with TonB-III (Noinaj et al., 2010). For TolQRA, force application may influence how TolA-III perturb an OM complex comprising the periplasmic protein TolB and the cell-wall binding lipoprotein Pal (Bonsor et al., 2009; Szczepaniak et al., 2020), leading to maintenance of OM lipid homeostasis, albeit via an elusive mechanism (Shrivastava et al., 2017; Tan & Chng, 2024). Indeed, the notable lack of structural information of these ‘stator-motor-effector’ complexes preclude detailed understanding of *pmf*-dependent force transduction across the cell envelope.

To bridge this gap, we hereby elucidate the cryo-electron microscopic (cryo-EM) structures of the *E. coli* TolQRA complex in detergent micelles, in distinct states at 3.6 Å and 4.2 Å resolutions. Our structures reveal how a single TolA N-terminal TM helix interacts with an asymmetric TolQ_5_R_2_ sub-complex at two different positions in the two states, which are in fact related by a 36° rotation of the TolQ pentamer. Combining further analysis of *in silico* predictions of the full TolQ_5_R_2_A complex, we introduce a mechanistic model that illustrates how a small rotary movement in the motor-stator complex can possibly be coupled to conformational changes in TolA-II to ultimately transmit a physical force for OM lipid homeostasis by the Tol-Pal complex.

## Results

### The inner membrane TolQRA complex can be stably purified, likely in a 5:2:1 stoichiometry

For structural characterization, we overexpressed *E. coli* TolQRA in BL21(λDE3) cells, and purified the complex using an octahistidine-tag at the C-terminus of TolR. The TolQRA complex was first extracted from total membrane fractions using the non-ionic detergent, n-dodecyl-β-D-maltoside (DDM), and subsequently replaced with (2,2-bis(3’-cyclohexylbutyl) propane-1,3-bis-β-D-maltopyranoside (CYMAL6-NG), which helped to enhance complex stability during cobalt(II) affinity purification. The purified complex eluted as a homogeneous peak on size-exclusion chromatography (SEC), with three distinct bands on SDS-PAGE corresponding to TolQ (25.6 kDa), His-TolR (15.4 kDa) and TolA (43.2 kDa) (**Fig. 1A**). Label-free tandem MS quantification estimated a stoichiometry of 4.5 TolQ: 1.5 TolR: 1 TolA; since homologous stator-motor complexes ExbBD and MotAB comprise 5 copies of ExbB/MotA and 2 copies of ExbD/MotB (Celia et al., 2019; Deme et al., 2020a; Santiveri et al., 2020), we conclude that we have successfully isolated the native TolQ_5_R_2_ subcomplex stably co-purified with one copy of TolA. On optimized cryo-EM grids, we observed homogenous particle distribution of our purified TolQ_5_R_2_A preparations, facilitating picking and sorting into well-resolved 2D classes of unique orientations (final ∼734K particles) (**Fig. 1B**, and **S1A**). Notably, the 5:2 ratio of TolQ:TolR was already discernible from specific ‘bottom’ orientations of the complex.

**Figure 1.**
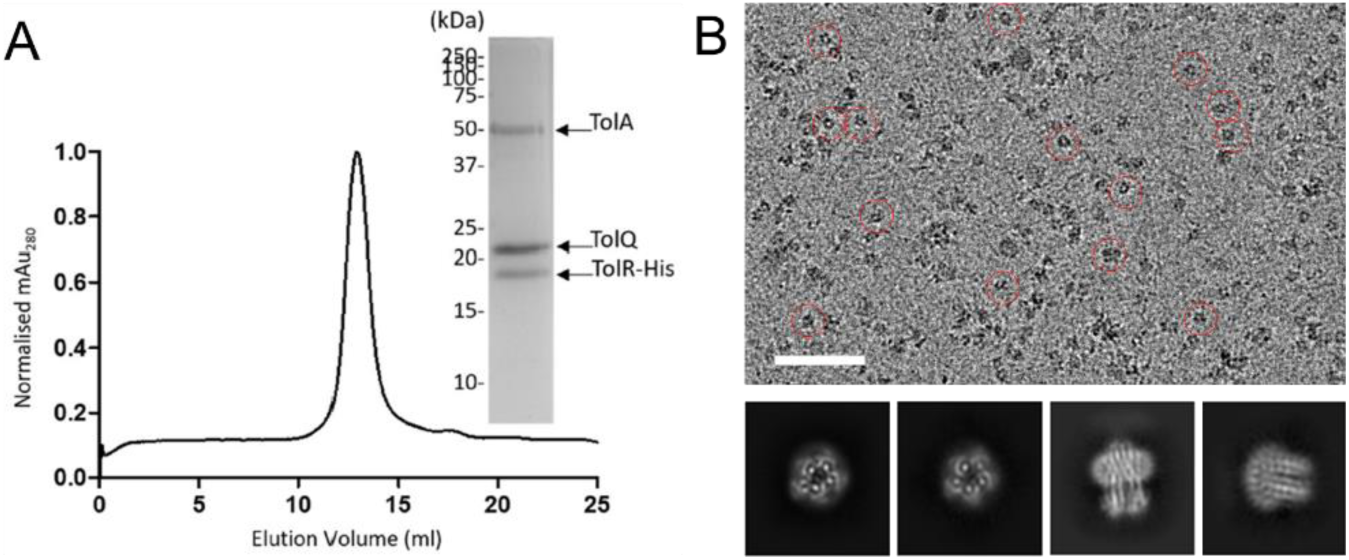
The *E. coli* TolQ_5_R_2_A complex can be purified to homogeneity. (**A**) Size exclusion chromatographic (SEC) and SDS-PAGE analyses of purified TolQ_5_R_2_A complexes in CYMAL6-NG detergent micelles. (**B**) *Top*: Representative cryo-EM image with several homogenous TolQ_5_R_2_A particles marked by *red* circles. Scale bar – 50 nm. *Bottom*: Representative 2D classes derived from 669,534 particles (using particle box size of 240 pixels (200 Å)) across 6,036 micrographs (see **Fig. S1**).

### The TolQ_5_R_2_A complex exists in two distinct states

Initial 3D reconstruction without application of symmetry operators afforded a density map at 3.6 Å nominal resolution (∼165K particles) (**Fig. S1B**). We fitted an AlphaFold3 predicted model (**Fig. S2**) of TolQ_5_R_2_A as rigid bodies into the density, immediately revealing that our initial map encompasses only the transmembrane and cytoplasmic regions of the complex. The periplasmic regions are likely highly flexible, and therefore largely averaged out in the final models, consistent with the observation of low-resolution densities above the main complex (**Fig. S1**).

Density for five protomers of TolQ (each comprising three TM helices) enclosing two N-terminal TM helices of TolR were clearly defined. In addition, the N-terminal TM helix of TolA (TolA-I) was fitted into density observed at the periphery of one TolQ subunit. Interestingly, we also detected extra density (at lower contour) at another TolQ protomer, suggesting the possibility of a second TolA TM helix (**Fig. S1**). Given that stoichiometric estimation indicated only one copy of TolA in the purified complex, we hypothesized that our initial map could reflect a mixture of the TolQ_5_R_2_A complex in two distinct states. Indeed, further 3D classification yielded two discrete volumes, map 1 (EMD-**62050**, 59% of particles) (**Fig. 2A**) and map 2 (EMD-**62251**, 41% of particles) (**Fig. 2C**), with the presumed TolA TM helix density at two separate TolQ protomers.

**Figure 2.**
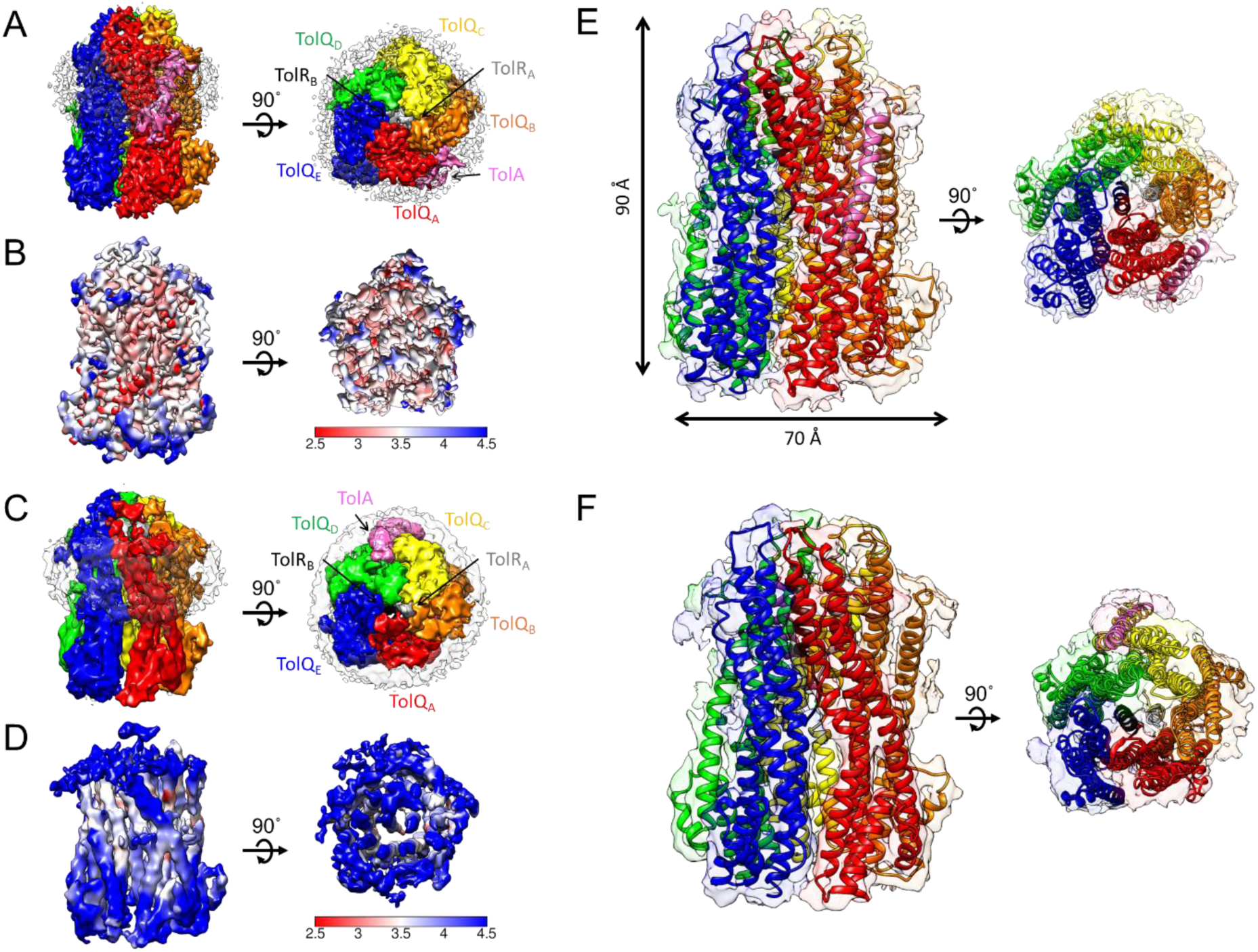
Purified TolQ_5_R_2_A complexes are captured in two distinct states in detergent micelles, where TolA interacts with a different TolQ protomer in each state. (**A, C**) Periplasmic and side views of the density maps (unsharpened; contour level of 0.06) of the transmembrane and cytoplasmic domains of TolQ_5_R_2_A in (**A**) state **A** (EMD-**62050**, 3.6 Å resolution) and (**C**) state **B** (EMD-**62251**, 4.3 Å resolution) with the protein surface densities colored *red*/*orange*/*yellow*/*green*/*blue* for TolQ_A-E_, *grey*/*black* for TolR_A-B_, *pink* for TolA, and *white* for detergent micelle (*transparency 80%*), respectively. (**B, D**) Sharpened local resolution maps were calculated by cryoSPARC (Punjani et al., 2017) and illustrated with pseudo-color representation of per-voxel resolution for both (**B**) state **A**, and (**D**) state **B**. **(E, F)** Cartoon illustrations of TolQ_5_R_2_A structures in (**E**) state **A** (PDB **9K49**) and (**F**) state **B** (PDB **9KCH**), showing well-fitted and refined backbones within sharpened protein surface densities (*transparency 80%*). For state **A**, side chains are also well-fitted and refined, but not shown.

The major TolQ_5_R_2_A state represented by density map 1 has 3.6 Å nominal resolution (3.14 – 4.14 Å, **Figs. 2B** and **S3A**). After rigid body fitting of the AF3 model and trimming of residues beyond the density, real-space refinement (including side chains) was performed to obtain a well-fitted structural model (state **A**, PDB **9K49**) (**Fig. 2E** and **S4**). On the other hand, the minor TolQ_5_R_2_A state represented by density map 2 has 4.2 Å nominal resolution (3.42 – 7.70 Å, **Figs. 2D** and **S3B**). Owing to its lower resolution, we only performed real-space refinement for the main chains, deriving the backbone structural model for state **B** (PDB **9KCH**) (**Fig. 2F** and **S5**).

### The structural model of the major TolQ_5_R_2_A state reveals the asymmetric architecture of the TolQR subcomplex

As with known structures of homologous motor-stator complexes such as ExbB_5_D_2_ and MotA_5_B_2_, five TolQ protomers form a pore that encompasses two TolR N-terminal helices in the major state **A** of the TolQ_5_R_2_A complex. For each TolQ subunit, almost the entire molecule was well modelled (D8-T223), featuring essentially a three-helix bundle that spans the lipid bilayer and extends deep (40 Å) into the cytoplasm (**Fig. 2E**). These protomers are nearly identical, aligning well in a pairwise fashion as rigid bodies (**Fig. S6**). In the TolQ_5_ pentameric structure, each protomer contributes two helices (TMH-2 and TMH-3) to line the central hydrophobic pore (housing the TolR dimer), which is connected to a hydrophilic vestibule formed by the cytoplasmic regions of the five TolQs. The interfaces between TMH-2 of one TolQ and TMH-3 of an adjacent TolQ are consistent with specific contacts previously suggested by suppressor analyses or engineered disulfide bonds (Goemaere et al., 2007; Zhang et al., 2011) (**Fig. S7**). Interestingly, we observed that each TolQ has a significantly different pseudo-radial tilt, with TolQ_A_ being most vertically positioned, and TolQ_C_/TolQ_D_, TolQ_B_ and TolQ_E_ tilted relatively ∼5°, ∼10° and ∼20°, respectively, overall bringing the periplasmic ends of the TolQ rigid protomers closer to each other in an asymmetric fashion (**Fig. 3A**).

**Figure 3.**
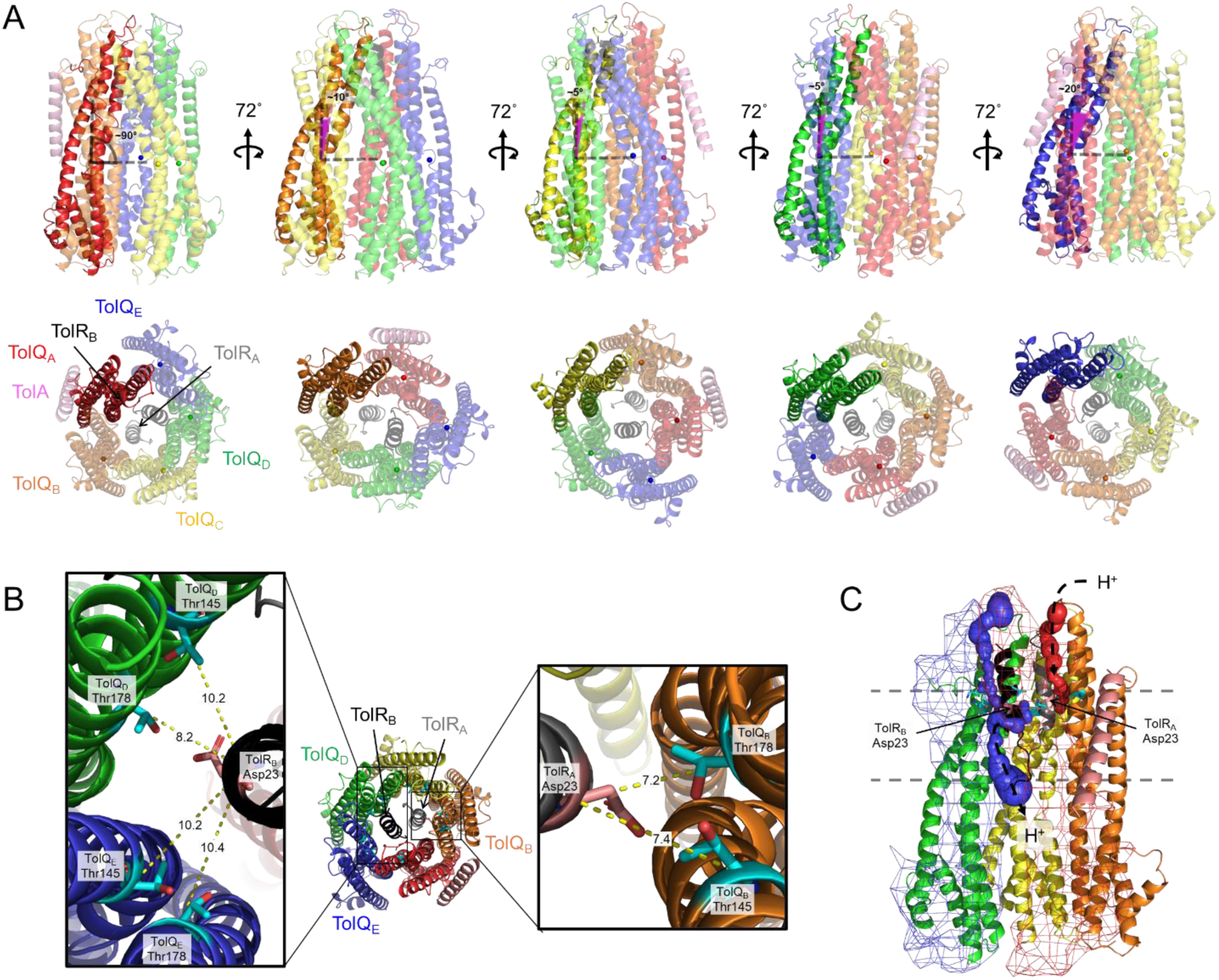
The architecture of the TolQ_5_R_2_ subcomplex reveals asymmetric spatial arrangements of TolQ protomers and TolR dimers, as illustrated in TolQ_5_R_2_A state **A** (PDB **9K49**). (**A**) Cartoon representations (side and top views) depicting the pseudo-radial tilts (*purple*) of TolQ_A_, and the other TolQs relative to TolQ_A,_ which were estimated by the angle TMH-1 of each TolQ made with the plane defined by the center-of-masses of all five TolQs. (**B**) Cartoon representations (top view) of the TolQ_5_R_2_A complex showing that Asp^23^ of TolR_A_ is positioned close to, and likely interacts with Thr^145^/Thr^178^ on TolQ_B_ alone (right inset), while Asp^23^ on TolR_B_ is positioned between TolQ_D_ and TolQ_E_, further from Thr^145^/Thr^178^ on both TolQs (left inset). Asp^23^ and Thr^145/178^ residues are represented by *cyan* and *pink sticks*, respectively, with oxygen heteroatoms colored *red*. C_α_-C_α_ distances (in Å) are indicated. (**C**) Cartoon representation (side view) of the TolQ_5_R_2_A complex illustrating accessibilities of Asp^23^ of TolR_A_ only to the periplasm, and of Asp^23^ of TolR_B_ to both the cytoplasm and periplasm, via channels revealed using MOLE 2.5 (Sehnal et al., 2013). The corresponding channels are represented as *red* and *blue spheres*, while TolQ_A_ and TolQ_E_ are represented in *red* and *blue mesh surfaces* (width, 2.5) for clarity. Inferred proton movement for protonation or deprotonation of the respective Asp^23^ residues are indicated.

The asymmetric tilting of TolQ protomers may be due to asymmetric interactions with the dimer of TolR N-terminal helices in the central hydrophobic pore, where alternating Thr^145^s (on TolQ TMH-2s) and Thr^178^s (on TolQ TMH-3s) form a ring of polar residues. Each TolR is clearly visible, with a short segment of unstructured tail (L12-V19) leading into a TM helix (P20-A34) that presents Asp^23^ for interaction with the TolQ Thr residues (**Fig. 3B**). Mutation suppressor analysis across the TolQ-TolR interface are thought to modulate the interactions between there three conserved residues, TolR-Asp^23^, TolQ-Thr^145^ and TolQ-Thr^178^, which are known to be essential for proton passage, and coupling *pmf* to (rotary) movement of the complex (Deme et al., 2020b; Goemaere et al., 2007; Zhang et al., 2009a) (**Fig. S7**). Interestingly, we were able to discern non-equivalent biochemical environments for the two TolR-Asp^23^s. For the first TolR (TolR_A_), Asp^23^ is closely juxtaposed to possibly interact well with both Thr^145^ (C_α_-C_α_ 7.4 Å) and Thr^178^ (C_α_-C_α_ 7.2 Å) on the same TolQ_B_ partner (**Fig. 3B inset**). On the other hand, Asp^23^ on the second TolR (TolR_B_) faces and resides between Thr^145^ of TolQ_E_ (C_α_-C_α_ 10.2 Å) and Thr^178^ of TolQ_D_ (C_α_-C_α_ 8.2 Å), albeit slightly further away and vertically displaced, suggesting less optimal interactions (**Fig. 3B inset**). It is possible that these two Asps have different protonation states in these two environments, which may ultimately influence the pseudo-radial tilts of the corresponding TolQ protomers, particularly TolQ_B_ and TolQ_E_. In fact, channel analysis of the TolQ_5_R_2_A complex revealed that Asp^23^ of TolR_A_ may only be accessible from the periplasm, while that of TolR_B_ is exposed to both the cytoplasm and periplasm (MOLE 2.5) (Sehnal et al., 2013) (**Fig. 3C**). In the context of the *pmf*, where proton concentrations are locally higher at the periplasmic side of the IM, we propose that TolR_A_ and TolR_B_ Asp^23^s are likely protonated and deprotonated, respectively, in the asymmetric TolQ_5_R_2_ subcomplex.

### TolA interacts in the same manner with different TolQ protomers in the major and minor states of the TolQ_5_R_2_A complex

In the major state **A** of the TolQ_5_R_2_A complex, TolA binds primarily to the TolQ_A_ protomer. Here, the TM of TolA (Q7-S33, TolA-I) is well modelled (**Fig. S4G**), and positioned obliquely (∼47°) to TMH-1 of its TolQ_A_ partner (**Fig. 4A**). The TolA-TolQ interface is extensive, and is centered near the inner leaflet of the IM, comprising a largely-hydrophobic TolQ_A/B_-facing residues (Lys^10^, Leu^11^, Ala^14^, Ile^15^, Ser^18^, His^22^, Phe^26^, Leu^29^, Ile^30^, Ser^32^, Ser^33^) on TolA and vice versa on TolQ_A_ TMH-1 (Leu^7^, Ile^25^, Ile^29^, Ala^33^, Ile^36^, Gln^37^, His^126^), and TolQ_B_ (Leu^15^, Leu^16^, Phe^143^, Trp^147^, His^151^). Ser^18^, His^22^, Leu^29^, and Ser^33^ of TolA forms the conserved SHLS motif that has previously been shown to be important for interactions with TolQ, and TolQ suppressors of TolA^S18L^ or TolA^H22R^ mutants are consistent with this observed TolA-TolQ interface (Germon et al., 1998, 2001; Karlsson et al., 1993; Koebnik et al., 1993; Zhang et al., 2009b) (**Fig. 4B**). For the minor state **B** of the TolQ_5_R_2_A complex, only the protein backbones could be confidently modelled (**Figs. 2C and S5**). We saw essentially the same asymmetric TolQ_5_R_2_ subcomplex, with the only difference being the TolA TM interacting now primarily with TolQ_C_ (**Fig. 2C**). Given the extensive interface, TolA likely stays with same TolQ protomer during rotary action (**Fig. S8**), and these two states we have captured may represent distinct rotational states, possibly correlated with alternating protonation/deprotonation of the pair of Asp^23^ residues in the TolR dimer (**Fig. 4C**).

**Figure 4.**
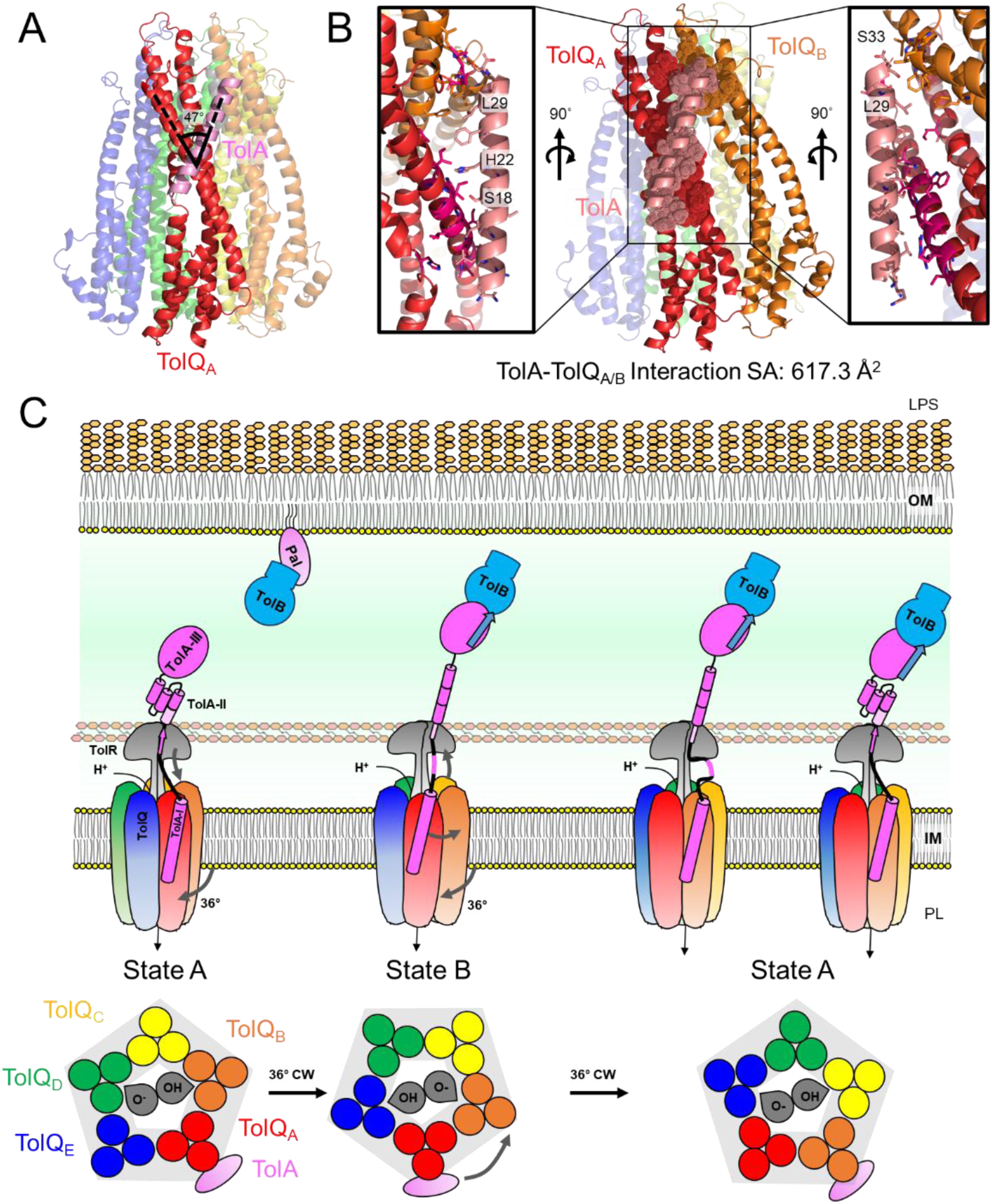
Proton translocation across the TolQ_5_R_2_A complex drives rotary motion of the stable motor-effector TolQ_5_TolA sub-complex, which may in turn induce conformational changes in TolA-II that facilitates extension across the cell envelope for interaction with TolB/Pal. (**A**) Cartoon representation (side view) depicting how the TolA TM helix is oriented relative to TolQ_A_ TMH-1. (**B**) Cartoon representations showing extensive interaction surface area (617.3 Å^2^, PISA) (Krissinel & Henrick, 2007) between TolA TM helix (TolA-I) and TolQ_A_/TolQ_B_. Specific residues in contact on TolA (*pink dots* and *sticks*), TolQ_A_ (*red dots* and *sticks*) and TolQ_B_ (*purple dots* and *sticks*) are shown in the respective insets. Highly conserved residues of the SHLS motif on TolA are indicated (Ashkenazy et al., 2016). (**C**) A proposed model for force transduction by the TolQ_5_R_2_A complex. Alternating protonation and deprotonation of Asp^23^ residues in the TolR_2_ stator powers a 36°-step rotation that switches the TolQ_5_R_2_A complex from state **A** to state **B**, which possibly transmits a pulling force on TolA-II, disrupting predicted interactions between TolA and the TolR periplasmic domains (*pink arrow*), and perhaps triggering re-organization of the TolA-II helical domain (*pink cylinders*) into an extended form for interaction with TolB/Pal across the periplasm. In the next 36°-step rotation of the TolQ_5_ motor, accompanied movement of TolA-I might be prevented due to the TolA-II domain, thus TolA-I become dislodged and re-positioned to the next TolQ protomer, effectively resetting the complex to state **A**. This reset allows reformation of the predicted TolA-TolR interactions, and possible re-bundling of the TolA-II helical domain, which thus represents a physical force pulling TolB towards the IM. O^-^, and OH in TolR represents Asp^23^ deprotonated and protonated states, respectively.

## Discussion

While it is widely accepted that proton translocation across the IM TolQRA complex is coupled to conformational changes in TolA to bind TolB across the cell envelope, how this force transduction occurs is mechanistically unclear. Here, to gain insights into TolQRA function, we have stably purified the TolQ_5_R_2_A complex from *E. coli*, and elucidated the cryo-EM structures of two distinct states in detergent micelles at 3.6 Å and 4.2 Å resolutions. Owing to the flexibility of the periplasmic regions of TolR and TolA, only the membrane and cytoplasmic regions of TolQ_5_R_2_A were well resolved. Our structures have revealed how the TolA TM helix interacts with two different TolQ protomers in the asymmetric TolQ_5_R_2_ sub-complex, which features a TolQ pentamer encapsulating a pair of TolR TM helices in non-identical biochemical environments. Our work lays the foundation for understanding how proton translocation may lead to rotary movement within the TolQ_5_R_2_A complex.

In *pmf*-utilizing motor-stator complexes, including MotA_5_B_2_, ExbB_5_D_2_, and TolQ_5_R_2_, the motor (MotA, ExbB, or TolQ) pentamer is believed to exhibit rotary motion around the stator (MotB, ExbD, or TolR) dimer, which is kept stationary via the binding of peptidoglycan through their periplasmic domains. Proton passage across the central pore of the motor presumably causes alternating protonation and deprotonation of a pair of Asps on the TM helices of the stator, influencing polar contacts that likely drives rotary movement (Deme et al., 2020b; Santiveri et al., 2020). In TolQ_5_R_2_A, since TolA interacts extensively with one TolQ protomer (**Fig. 4B**), its movement should be tied to this interface in the absence of other external forces; as the TolQ pentamer rotates ten steps of 36° around the TolR dimer, one can expect five unique rotary states, where TolA would be found at each TolQ position in the asymmetric TolQ_5_R_2_ subcomplex (**Fig. S8**, states **A/A’** to **E/E’**). If TolQ_5_R_2_A action requires full cycle rotation, these five states would be expected to be energetically comparable for smooth rotary motion, and thus equally populated in an ensemble. Interestingly, however, our structural analysis has only captured two stable states (**A** and **B**), which appear in fact to be related by just a 36° turn (**Figs. 4C** and **S8**). This observation suggests that TolQ_5_R_2_A might not perform a full rotation during force transduction, but undergoes some form of reset, perhaps via repositioning of TolA, to “return” from one state to another beyond that one small turn.

To facilitate this reset, there may be an opposing force to prevent the TM helix of TolA from moving with further rotary action of the TolQ pentamer. We speculate that this reset may depend on the periplasmic regions of TolR and TolA. The IM proximal periplasmic domain of TolA (i.e. TolA-II) possesses heptad repeats and is believed to adopt largely α-helical structures (Levengood et al., 1991; Witty, 2002), which is consistent with more recent AlphaFold2 predictions (Jumper et al., 2021) (**Fig. S9**). Interestingly, the AlphaFold3 model of the *E. coli* TolQ_5_R_2_A complex also revealed a possible interaction, in the form of β-strand augmentation, between the TolR periplasmic domains and a highly conserved yet unstructured region linking TolA-I and TolA-II (**Fig. S2, inset**) (Abramson et al., 2024). We posit that rotation of the TolQ pentamer during the transition from state **A** to state **B** in the 36° turn may exert a force upon this unstructured region and disrupt its interaction with TolR, in turn pulling on and affecting the α-helical periplasmic TolA-II domain (**Fig. 4C**). However, further rotation beyond state **B** could be prevented by the rest of TolA-II – as the TolQ pentamer rotates another 36°, the TolA TM helix may be dislodged and re-positioned to the next protomer, effectively returning it to state **A** (reset).

How then does this turn-and-reset mechanism translate to trans-envelope force transduction? Despite being the least well conserved protein in the Tol-Pal complex, AlphaFold2 models of TolA from different organisms still reveal long α-helical segments within TolA-II, but with varying positions for helical breaks (**Fig. S9**). Solution X-ray analyses of TolA in *Pseudomonas aeruginosa* have also hinted at TolA-II structure possibly more consistent with helical hairpins or bundles (Witty, 2002). Overall, these observations suggest that TolA-II may be α-helical but could adopt different states (from a single long helix to higher-ordered structures). While the exact mechanisms will require further investigation, one model for how TolQ rotary movement transduces a force across the cell envelope may be via conformational changes in TolA-II (**Fig. 4C**). Assuming TolA-II initially exists as a stable helical bundle, rotary movement-induced pulling on TolA could trigger unbundling, allowing TolA-II to now adopt a single extended helix to reach out to OM partners (e.g. TolB). As the rotary state resets, the pulling force on TolA is alleviated, allowing TolA-TolR β-augmentation to re-establish, and possibly facilitating re-formation of the helical bundle. Such re-organization of extended TolA-II into a higher-ordered structure may thus exert a force across the cell envelope, activating processes that need to happen near the OM. TolA is believed to interact with and tug on TolB at the OM; doing so disrupts the latter’s interaction with Pal, which can modulate Pal’s accessibility to cell wall (Szczepaniak et al., 2020). However, how such force transduction across the cell envelope eventually enables the Tol-Pal complex to execute its primary role in maintaining OM lipid homeostasis remains to be investigated (Shrivastava et al., 2017; Tan & Chng, 2024).

In a recent parallel study, the high resolution cryo-EM structures of the TolQ_5_R_2_A and ExbB_5_D_2_-TonB complexes have been determined (Celia et al., 2024). Similar to our findings here, two positions for the TolA TM helix were observed in their TolQ_5_R_2_A_(2)_ structure, though unresolved and presented as one state. Interestingly, three rotary states for ExbB_5_D_2_-TonB were captured, which apparently represented states **A-C** in the rotation cycle (**Fig. S8**), suggesting that force transduction in this homologous complex may require two 36^°^ turns instead of one, before reset. This may be unsurprising given the completely different sequence and expected (lack of) structure of the proline-rich TonB-II domain, which will influence how rotary force will be translated into a physical force across the cell envelope. Even so, a model alternative to the turn-and-reset mechanism was proposed in the study, where full 360^°^ rotation(s) of ExbB_5_-TonB reels in the largely unstructured TonB-II domain around the ExbD_2_ stator for force transduction. It is possible that TolQ_5_R_2_A action follows such a model; however, the return/reset of the system to its initial state (i.e. unreeling of TonB-II/TolA-II) might be energetically challenging.

## Acknowledgements

J.Y., C.G.C., and N.Z.-L.L. were supported by the National University of Singapore Graduate School of Integrative Sciences and Engineering Scholarship (ISEP). This work was supported by the Singapore Ministry of Education Academic Research Fund Tier 1 grant (National University of Singapore Faculty of Science Preparatory Grant Scheme) and the Singapore Ministry of Health National Medical Research Council under its Cooperative Basic Research Grant (NMRC/CBRG/0072/2014) (all to S.-S.C.). The authors would also like to acknowledge Dr Jian SHI from the Centre for Bio-Imaging Sciences (CBIS) at the National University of Singapore (NUS) for micrograph collection and microscope facility support.

## Data Availability

3D cryo-EM maps of two TolQ_5_R_2_A states have been deposited in the Electron Microscopy Data Bank under accession numbers EMD-**62050** (state **A**), EMD-**62251** (state **B**). Two atomic coordinate files have also been deposited in the Protein Data Bank under the accession numbers **9K49** (state **A**) and **9KCH** (state **B**).

## Competing Interest Statement

The authors declare no competing interests.

## Methods

### Bacterial strains and plasmids

All strains, plasmids, and primers used are listed in **Supplementary Table 1, 2, and 3** respectively.

### Growth conditions

Lysogeny Broth (LB) and agar were prepared at 2.5% (w/v), and 1% (w/v) respectively. Unless otherwise noted, ampicillin (Amp) (Sigma-Aldrich, MO, USA) was used at a concentration of 200 μg/mL.

### Over-expression and purification of TolQRA complex

To perform structural analysis, TolQR(-His)A protein complexes were over-expressed and purified from BL21(λDE3) cells transformed with pET22/42*tolQR(-His)A*. An overnight 5-mL culture was grown from a single colony in LB broth supplemented with appropriate antibiotics at 37 °C. The overnight cell culture was then used to inoculate a 250-mL culture and grown at the same temperature until OD_600_ reached ∼ 0.7. This culture was then further used to inoculate ten 1.5-L cultures and grown at the same temperature until the OD_600_ has again reached ∼ 0.7. For induction, 1.0 mM of isopropyl β-D-1-thiogalactopyranoside (IPTG) (Axil Scientific, Singapore) was added and the culture was grown for another 3 h at 37 °C. Cells were harvested by centrifugation at 4,700 x *g* for 20 min and stored at -80 °C. Cell pellet was resuspended then resuspended in phosphate-buffer saline (PBS) containing 1 mM phenylmethylsulfonyl fluoride (PMSF) (Calbiochem), 50 μg/mL DNase I (Sigma-Aldrich), and 100 μg/mL lysozyme (Calbiochem). Cells were lysed with two passes at high pressure in a French Press G-M® High Pressure Cell Disruptor (20,000 psi, Glen Mills). The lysed sample was centrifuged (4700 rpm, 4 °C, 20 mins) to remove cell debris. Ultracentrifugation (36,000 rpm, 4 °C, 1 h) was used to separate the membrane and soluble fractions in the supernatant. The pellet containing the membrane fraction was resuspended in membrane solubilising buffer (PBS; 20 mM NaH_2_PO_4_/Na_2_HPO_4_ pH 7.5, 300 mM NaCl, 20 mM imidazole, and 1 % n-dodecyl β-D-maltoside (DDM, Calbiochem)) and the resuspension was incubated at 4°C for 2 h with agitation. DDM-solubilised membrane fraction was subjected to another round of ultracentrifugation (36,000 rpm, 4 °C, 1 h) to isolate the extracted membrane proteins. The supernatant containing the solubilised membrane fraction was then incubated with 1 mL Co-NTA cobalt resin (TALON® Metal Affinity Resin, Takara Bio Inc, Shiga, Japan), pre-equilibrated with 20 mL of wash buffer (PBS containing 0.04 mM CYMAL6-NG and 20 mM imidazole) in a column for 3 h at 4 °C with rocking. The mixture was allowed to drain by gravity, and the flowthrough was passed through the resin bed twice, before washing vigorously with 10 x 10 mL of wash buffer and eluted with 10 mL of elution buffer (PBS containing 0.04 mM CYMAL6-NG and 200 mM imidazole). The eluate was concentrated in an Amicon Ultra 100 kDa cut-off ultra-filtration device (Merck Millipore) by centrifugation at 4,000 x *g* to ∼500 μL. Proteins were further purified by size-exclusion chromatography (AKTA Pure, GE Healthcare, UK) at 4 °C on a prepacked Superose 6 increase 10/300 GL column, using PBS containing 0.04 mM CYMAL6-NG as the eluent.

### SDS-PAGE

All samples subjected to SDS-PAGE were mixed 1:1 with 2X Laemmli buffer. The samples were subjected to boiling at 100 °C for 10 min. Equal volumes of the samples were loaded onto the gels. As indicated in the figure legends, SDS-PAGE was performed using either 12% Tris.HCl gels at 200 V for 45 min. After SDS-PAGE, gels were visualized using Coomassie Blue staining by *G*:Box Chemi-XT4 (Genesys version 1.4.3.0, Syngene).

### Cryo-EM grid preparation and data acquisition

For sample preparation, 3.0 μL of the protein sample at a concentration of 15 mg/mL was applied to glow-discharged Quantifoil holey carbon grids (1.2/1.3, 200 mesh, Electron Microscopy Sciences). Grids were blotted for 3 s with 100% relative humidity at 4 °C and plunge-frozen in liquid ethane cooled by liquid nitrogen using a Vitrobot System (Gatan). Cryo-EM data were collected at liquid nitrogen temperature on a Titan Krios electron microscope (Thermo Fisher Scientific), equipped with a K3 Summit direct electron detector (Gatan) and GIF Quantum energy filter. All cryo-EM movies were recorded in counting mode with SerialEM4 ^17^ with a slit width of 20 eV from the energy filter. Movies were acquired at nominal magnifications of 105,000 x, corresponding to a calibrated pixel size of 0.834 Å on the specimen level. The total exposure time of each movie was 5.2 s, resulting in a total dose of 56 electrons per Å^2^, fractionated into 40 frames. More details of electron microscopy data collection parameters are listed in **Supplementary Table 4**.

### Electron microscope image processing

cryoSPARC 4.6.0 (Punjani et al., 2017) was used to process the EM data according to the flowchart in **Supplementary Figure 1**. Dose-fractionated movies were corrected for motion using Patch Motion Correction. To obtain a sum of all frames for each movie, a dose-weighting scheme was applied, and this sum was used for all image-processing steps except for defocus determination. Defocus values of the summed images from all movie frames were calculated using patch CTF estimation without dose weighting. CTF-estimated micrographs were then denoised using a greyscale normalisation factor of 0.8, with an increased number of data pass-throughs during training to 400. Particle picking was performed using the blob picker followed by the template picker. Two- and three-dimensional (2D and 3D) classifications were carried out using "2D classification", "Ab-initio Reconstruction", and "Heterogeneous Refinement". 3D refinements were conducted using "Homogeneous Refinement" and "Non-Uniform Refinement". Reference motion correction was applied to the micrographs and particles using locally-refined 3D volumes obtained. Eventually, we obtained a cryo-EM map representing compositional heterogeneities of TolA in TolQ_5_R_2_A complexes. A 3D-classification separated the compositional heterogeneities into two states; state **A** (EMD-**62050**) and state **B** (EMD**-62251**). The overall resolutions for these maps were estimated based on the gold-standard criterion of Fourier shell correlation (FSC) = 0.143, while local resolution was estimated with "Local Resolution Estimation".

### Model building and refinement

An initial model of TolQ_5_R_2_A complex was built using AlphaFold3 (Abramson et al., 2024). The model was rigid-body fitted into the highest resolution cryo-EM map, state **A** (EMD-**62050** – 3.60 Å), and trimmed in Chimera (Pettersen et al., 2004), followed by density-fitting in COOT 0.9.8 (Emsley & Cowtan, 2004). Refinement in RealSpace using the program of Phenix 1.20.1-4487 (Adams et al., 2010) with default parameters yielded the TolQ_5_R_2_A structure for state **A** (PDB **9K49**). Next, the mainchain of the TolQ_5_R_2_A structure for state **A** was used as the initial model for density-fitting and refinement in the cryo-EM map of state **B** (EMD-**62251** – 4.19 Å), in COOT 0.9.8 (Emsley & Cowtan, 2004) and Phenix 1.20.1-4487 (Adams et al., 2010), to yield coordinates for the TolQ_5_R_2_A structure for state **B** (PDB **9KCH**). The final coordinates of the asymmetric units were checked using MolProbity (Davis et al., 2007). Maps and structures shown in the figures were generated using UCSF Chimera and COOT 0.9.8. The model building and refinement statistics are shown in Supplementary **Table 4**.

### Measurements and figure preparation

Measurements were calculated and figures were generated using Chimera (Pettersen et al., 2004), PyMOL, MOLE2.5 (Sehnal et al., 2013), and PISA (Krissinel & Henrick, 2007).

## Notes

### Competing Interest Statement

The authors have declared no competing interest.

