## Supporting Information for "Structural insights into the force-transducing mechanism of a motor-stator complex important for bacterial outer membrane lipid homeostasis"

**Affiliation:**

**This PDF file includes:**

Supplementary Figures 1 to 9

Supplementary Tables 1 to 4

Supplementary References

### Supplementary Figures

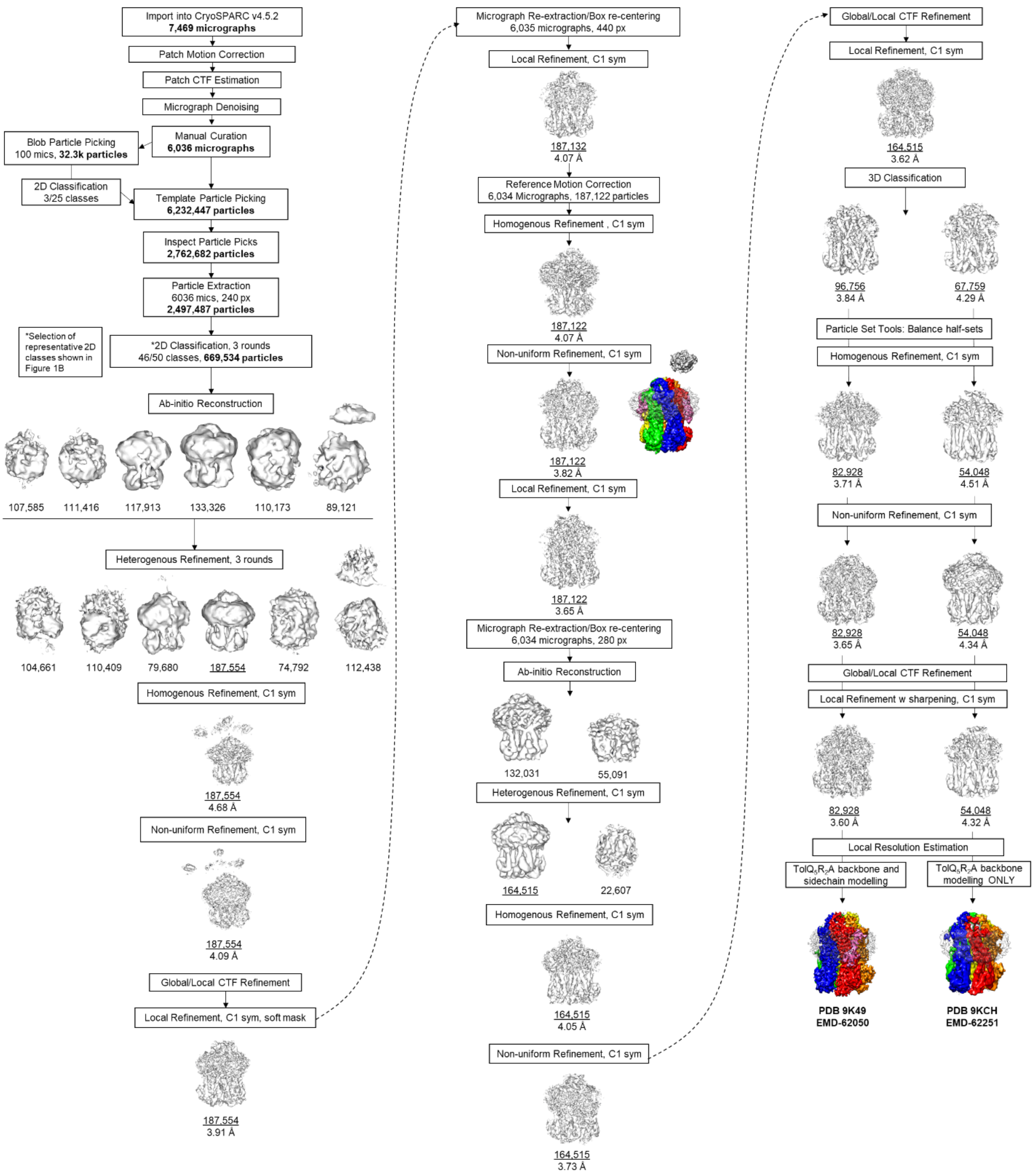

**Supplementary Figure 1**. Single-particle cryo-EM analysis of detergent-embedded $TolQ_5R_2A$ complexes. Data processing flowchart yielding the final maps (EMD**-62050** and EMD**-62251**) of two states representing spatial heterogeneity of TolA in $TolQ_5R_2A$. The structure (backbones and side chains) for state **A** of $TolQ_5R_2A$ (PDB **9K49**) was built and refined in EMD**-62050**, while that (backbones) of state **B** (PDB **9KCH**) was modelled in EMD**-62251**. Additional refinement details can be found in **Supplementary Table 4**.

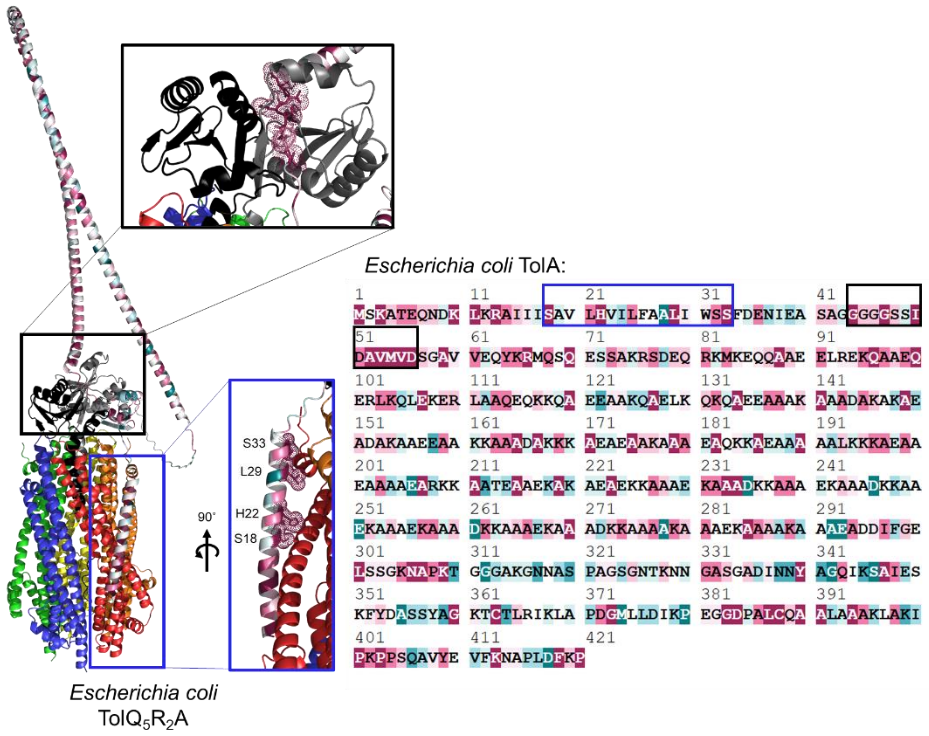

**Supplementary Figure 2**. The *in silico* model of the *E. coli* $TolQ_5R_2A$ complex hints at possible interactions between the TolR dimer and TolA in the periplasm. (**A**) Cartoon representation of the AlphaFold3 (Abramson et al., 2024) model of the full length $TolQ_5R_2A$ complex illustrating predicted interactions between $TolQ_A$ and TolA-I (*blue* inset), and between the periplasmic region of the TolR dimer and a highly-conserved unstructured region linking TolA-I and TolA-II (*black* inset). $TolQ_{A\text{-}E}$ are colored *red*/*orange*/*yellow*/*green*/*blue* and $TolR_{A/B}$ *grey*/*black*, respectively. Residues of TolA were colored according to their degree of conservation shown on the sequence on the *right*, ranging from highly-conserved residues in *maroon*, to *white*, and to poorly-conserved residues in *cyan* (Ashkenazy et al., 2016).

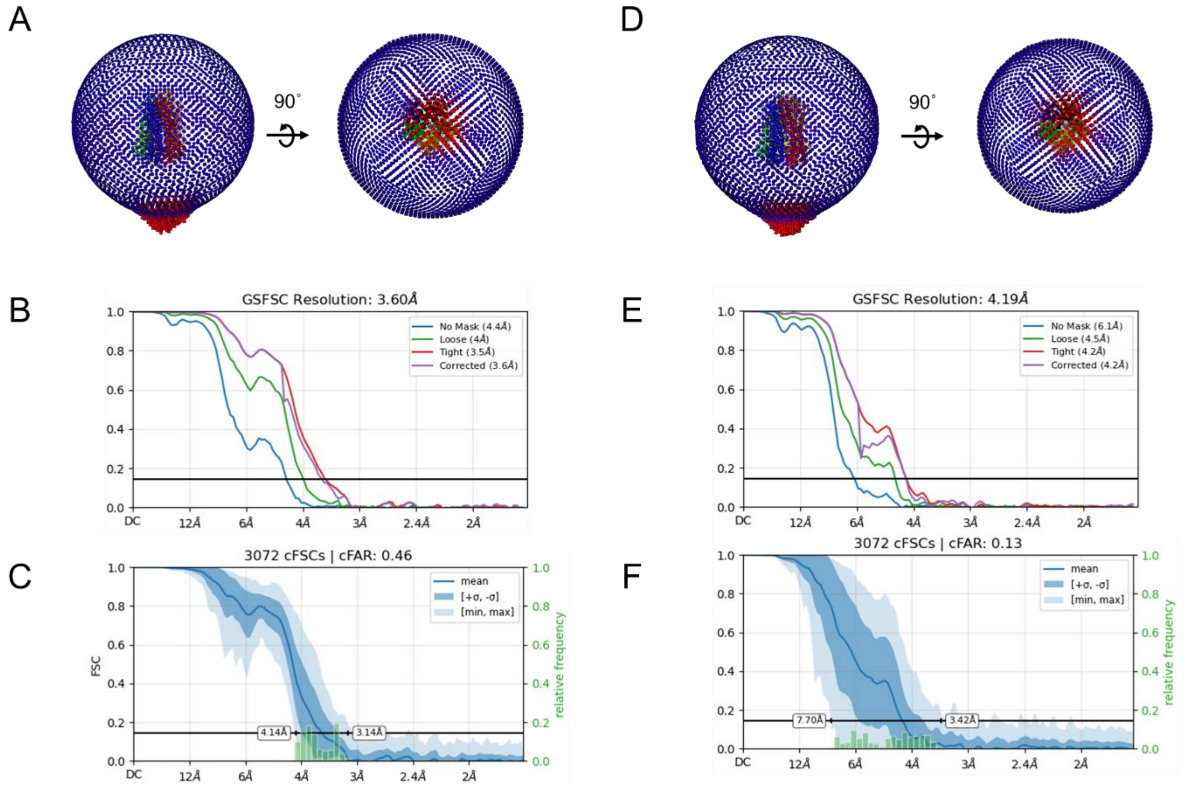

**Supplementary Figure 3.** Single-particle cryo-EM analysis of detergent-embedded $TolQ_5R_2A$ complexes. Relevant parameters (Euler angle distributions, GS-FSC plots, resolution distributions) for (**A-C**) state **A**, and (**D-F**) state **B** are presented. For Euler angle distributions, the height and color of each rod is proportional to the amount of particles visualized from the same specific orientation. The gold standard-Fourier Shell Correlation (GS-FSC) plots of unmasked and masked (loose, tight, and corrected) maps are derived from cryoSPARC (Punjani et al., 2017).

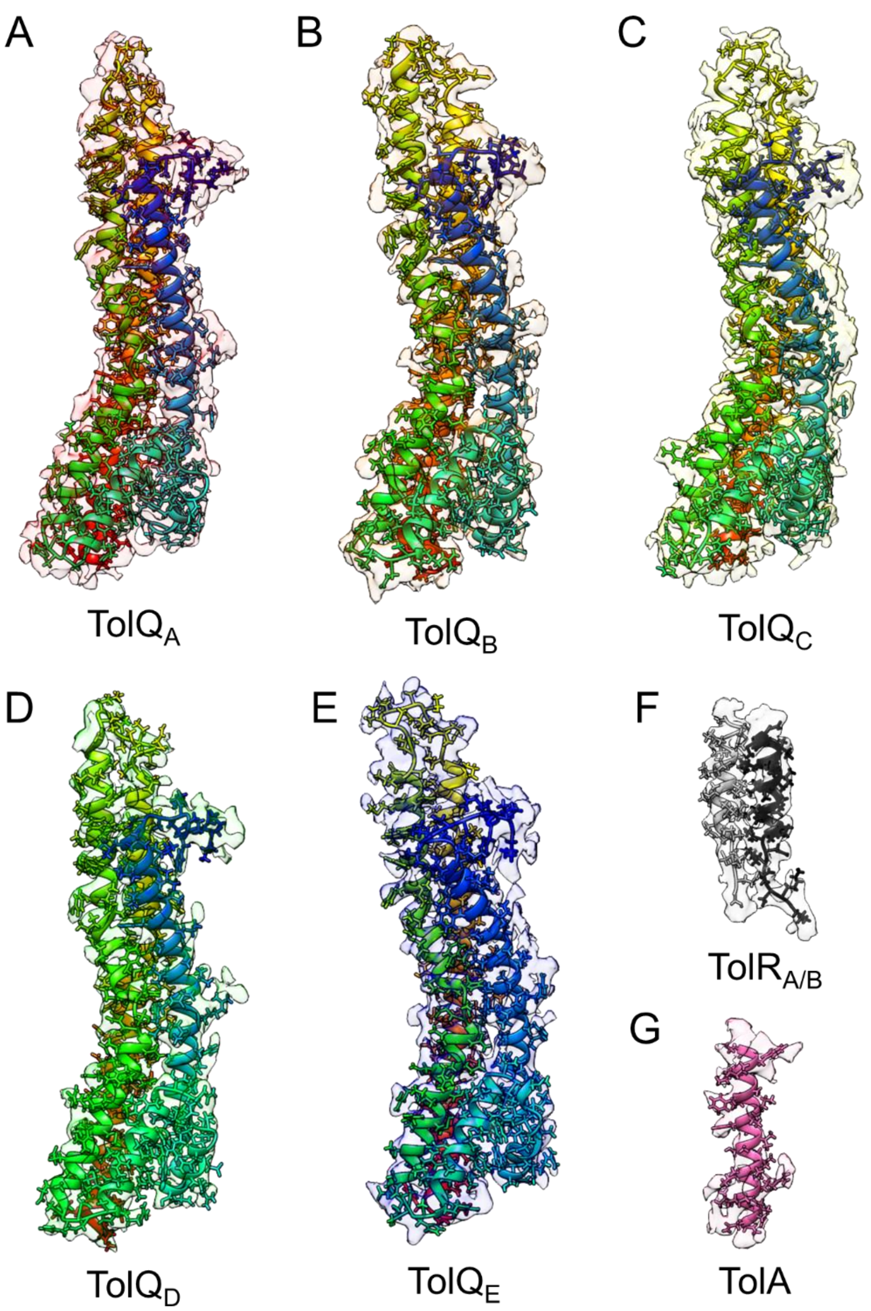

**Supplementary Figure 4**. Backbones and side chains of individual chains of the $TolQ_5R_2A$ complex (PDB **9K49**) are well-fitted into their corresponding densities in the map of state **A** (EMD-**62050**), *transparency 80%*, contour level of 0.06). (**A**) TolQ$_A$, (**B**) TolQ$_B$, (**C**) TolQ$_C$, (**D**) TolQ$_D$, (**E**) TolQ$_E$, (**F**) TolR$_{A/B}$, and (**G**) TolA.

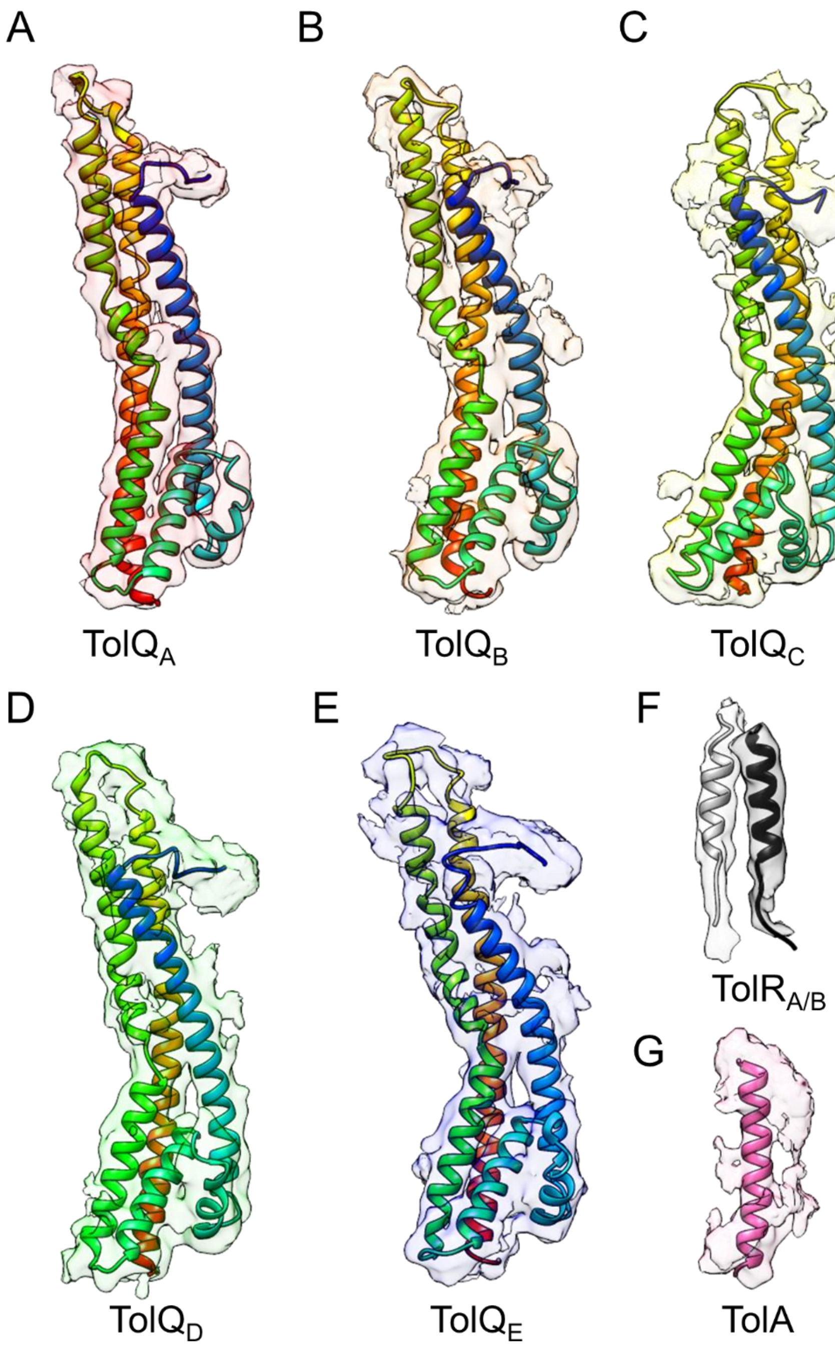
A
B
C
TolQA
TolQB
TolQC
D
E
F
TolRA/B
G
TolQD
TolQE
TolA

**Supplementary Figure 5**. Backbones of individual chains of the $TolQ_5R_2A$ complex (PDB **9KCH**) are well-fitted into their corresponding densities in the map of state **A** (EMD**-62251**, *transparency 80%*, contour level of 0.06). (**A**) $TolQ_A$, (**B**) $TolQ_B$, (**C**) $TolQ_C$, (**D**) $TolQ_D$, (**E**) $TolQ_E$, (**F**) $TolR_{A/B}$, and (**G**) TolA.

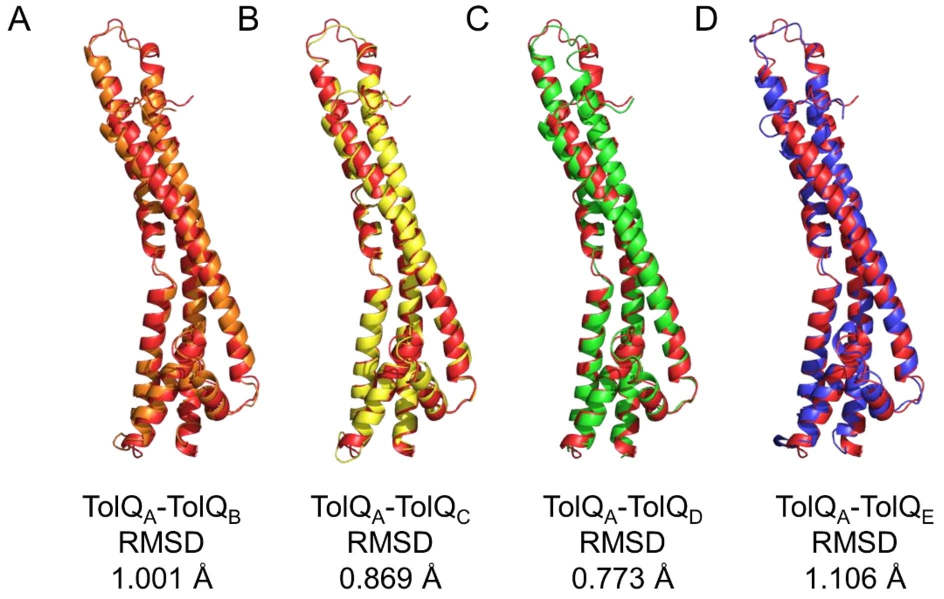

**Supplementary Figure 6**. TolQ protomers in the $TolQ_5R_2A$ complex are essentially identical. Pair-wise overlay of $TolQ_A$ (*red*) with $TolQ_{B/C/D/E}$ (*orange*/*yellow*/*green*/*blue*), revealing insignificant root mean square deviations (r.m.s.d. < 1 Å).

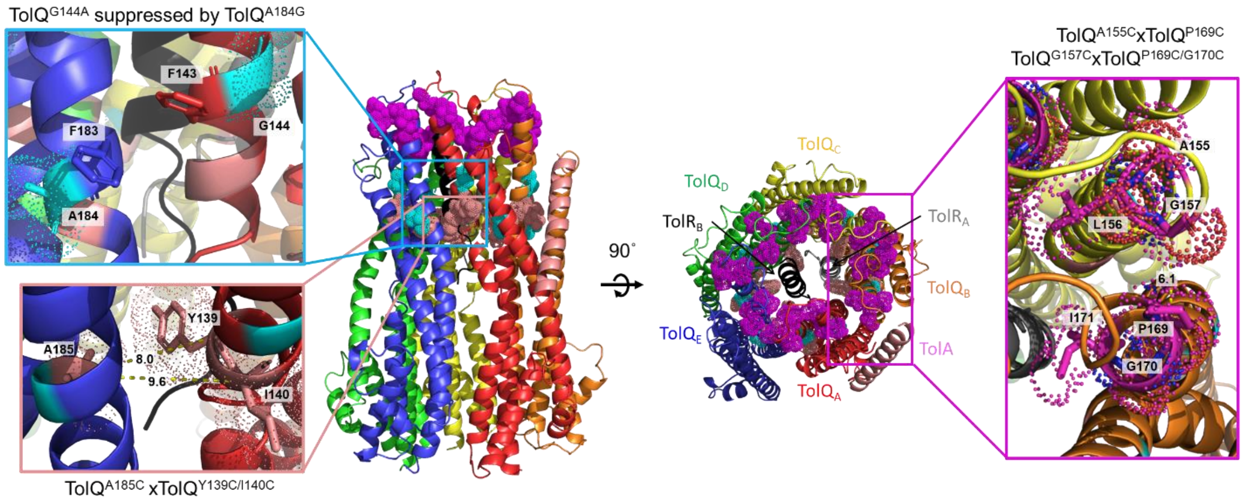

**Supplementary Figure 7**. Existing suppressor and disulfide bond analyses (Goemaere et al., 2007; Zhang et al., 2011) are in agreement with the architectural organization of the $TolQ_5R_2A$ complex. Cartoon representations of the $TolQ_5R_2A$ complex in state **A** (PDB **9K49**) showing residues previously demonstrated to exhibit genetic interactions, or facilitate disulfide bond formation when replaced with cysteines. In the *cyan* inset, TolQ$^{G144A}$ has been shown to suppress the TolQ$^{A184G}$ mutation (both represented by *cyan sticks* and *dots*), possibly by influencing orientations of, and interactions between adjacent Phe$^{143}$ and Phe$^{183}$ (*sticks*) across the TolQ-TolQ interface. In the *pink* and *purple* insets, TolQ$^{A185C}$ can be disulfide-bonded to TolQ$^{Y139C/I140C}$ (represented by *salmon sticks* and *dots*), and TolQ$^{A155C/G157C}$ to TolQ$^{P169C/G170C}$ (represented by *purple sticks* and *dots*) across the TolQ-TolQ interfaces. $C_\alpha$-$C_\alpha$ distances between relevant residues are indicated in Å.

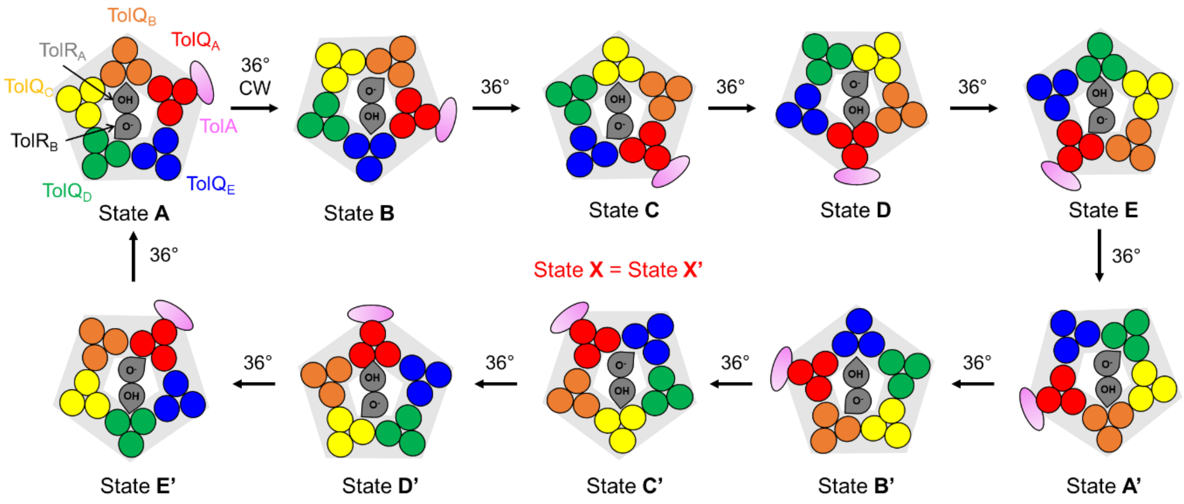

**Supplementary Figure 8**. In $TolQ_5R_2A$, ten steps of 36° rotation can theoretically occur in the $TolQ_5A$ subcomplex, in conjunction with alternating protonation (denoted OH) and deprotonation (denoted $O^-$) of the pair of $Asp^{23}$s in the $TolR_2$ dimer. A full rotation will give rise to five unique states (since state **X** = state **X'**) with TolA at each TolQ position in the asymmetric $TolQ_5R_2$ subcomplex. Of these, only two states (**A/A'** (PDB **9K49**) and **B/B'** (PDB **9KCH**)) have been observed. Note that the coloring of TolQ protomers in all states in this figure follows state **A**.

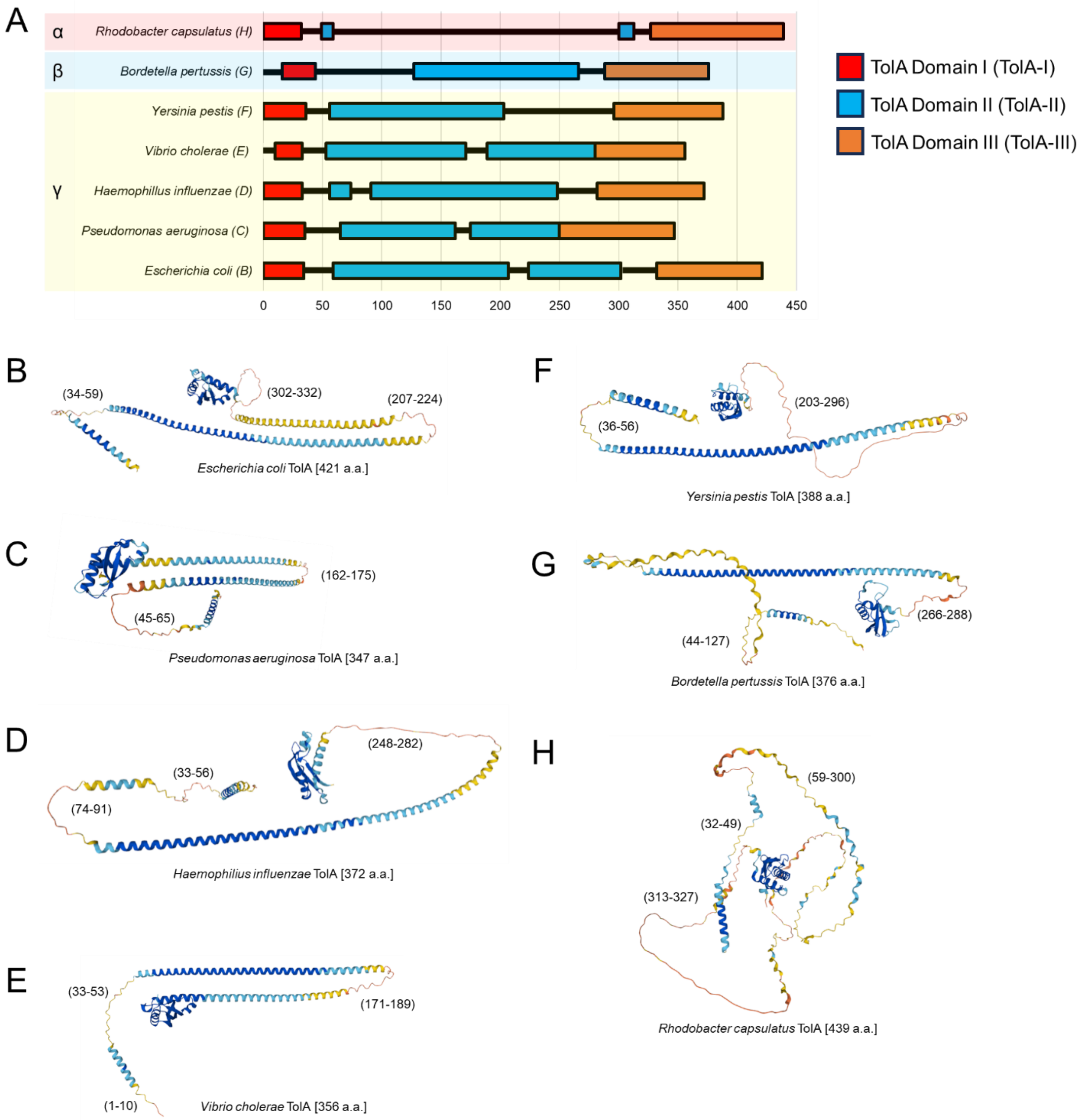

**Supplementary Figure 9**. Periplasmic TolA-II domain from different organisms adopt largely α-helical structures. (**A**) Predicted domain organizations of TolA from indicated organisms. Structured regions of domains I, II, and III, are represented by *red, blue*, and *orange bars*, respectively. The horizontal axis represents the number of amino acids. Classes of proteobacteria are indicated in colored shades. (**B-H**) AlphaFold2 (Jumper et al., 2021) models of TolA from different Gram-negative bacterial organisms reveal long α-helical segments within TolA-II, albeit with inconsistent predicted helical breaks (Sturgis, 2001). Cartoon representations are colored according to the confidence levels of prediction, from most confident (*dark blue*) to least confident (*yellow*). Numbers in parenthesis indicate the numbering of amino acids of unstructured regions of TolA.

### Supplementary Tables

**Supplementary Table 1**. Bacterial strains used in this study

| Strains | Relevant genotypes and characteristics | References |
|---|---|---|
| NovaBlue | *endA1 hsdR17 (rK12– mK12+) supE44 thi-1 recA1 gyrA96 relA1 lac F' proA+ B+ lacIq ZΔM15*::*Tn10* | Novagen |
| BL21(λDE3) | *fhuA2 lon ompT gal (λDE3) dcm* Δ*hsdS λDE3 = λ sBamHIo* Δ*EcoRI-B int*::*(lacI*::*PlacUV5*::*T7 gene1) i21* Δ*nin5* | Novagen |
| CG032 | BL21(λDE3) pET22/42*tolQR(-His)A* | This study |

**Supplementary Table 2.** Plasmids used in this study

| Plasmids | Relevant genotypes and characteristics | References |
| --- | --- | --- |
| pET22/42 | pT7($lac_{O}$) inducible expression vector, contains multiple cloning site of pET42a(+) in pET22b(+) backbone; $Amp^{R}$ | (Wu et al., 2006) |
| pET22/42*tolQR(-His)A* | Encodes full length TolQ, TolR with C-terminal His8 tag, and TolA; $Amp^{R}$ | This study |

**Supplementary Table 3.** Primers used in this study.

| Primers | Sequence (5' to 3') |
|---|---|
| TolQ-NdeI-FP | ATCGCATATGACTGACATGAATATCCTTGATTTGTTC |
| TolA-AvrII-RP | ATCGCCTAGGTTACGGTTTGAAGTCCAATGGC |
| TolR-SDM-8His-FP | CACCATCATCATCACCACCATCACTAAACATCTGCGTTTCCCTTGC |
| TolR-SDM-8His-RP | GTGATGATGATGGTGCTCGAGGATAGGCTGCGTCATTAAACCAAC |

**Supplementary Table 4.** Cryo-EM data collection, refinement and validation statistics

| | $TolQ_5R_2A$ State **A** (EMD-**62050**) (PDB **9K49**) | $TolQ_5R_2A$ State **B** (EMD-**62251**) (PDB **9KCH**) |
|---|---|---|
| **Data collection and processing** | | |
| Magnification | 105,000x | |
| Voltage (kV) | 300 | |
| Electron exposure (e–/Å$^2$) | 56 | |
| Defocus range (μm) | -0.5 to -2.0 | |
| Pixel size (Å) | 0.834 | |
| Symmetry imposed | C1 | |
| Initial particle images (no.) | 2,497,487 | |
| Final particle images (no.) | 82,928 | 54,048 |
| Map resolution (Å) | 3.60 (0.143) | 4.19 (0.143) |
| FSC threshold | | |
| Map resolution range (Å) | 3.14-4.14 | 3.42-7.70 |
| **Refinement** | State **A** | State **B** |
| Initial model used (PDB code) | AF3 | 9K49 |
| Model resolution (Å) | - | - |
| FSC threshold | | |
| Model resolution range (Å) | - | - |
| Map sharpening *B* factor (Å$^2$) | 116.6 | 124.3 |
| Composition | | |
| Chains | 8 | 8 |
| Protein residues | 1,161 | 1,161 |
| Non-hydrogen atoms | 9,121 | 9,121 |
| Ligands | 0 | 0 |
| R.m.s. deviations | | |
| Bond lengths (Å) | 0.004 | 0.004 |
| Bond angles (°) | 0.707 | 0.794 |
| Validation | | |
| MolProbity score | 1.81 | 1.73 |
| Clashscore | 5.03 | 7.25 |
| Poor rotamers (%) | 1.90 | 0.11 |
| Ramachandran plot | | |
| Favored (%) | 95.2 | 95.28 |
| Allowed (%) | 4.80 | 4.54 |
| Disallowed (%) | 0.00 | 0.17 |
